# Deficiency in Bhlhe40 impairs resistance to *H. polygyrus bakeri* and reveals novel Csf2rb-dependent regulation of anti-helminth immunity

**DOI:** 10.1101/687541

**Authors:** Nicholas N. Jarjour, Tara R. Bradstreet, Elizabeth A. Schwarzkopf, Melissa E. Cook, Chin-Wen Lai, Stanley Ching-Cheng Huang, Reshma Taneja, Thaddeus S. Stappenbeck, Steven J. Van Dyken, Joseph F. Urban, Brian T. Edelson

## Abstract

The cytokines GM-CSF and IL-5 are thought to possess largely divergent functions despite a shared dependence on the common beta (β_C_) chain to initiate signaling. Although IL-5 is part of the core type 2 cytokine signature and is required for protection against some helminths, it is dispensable for immunity to others, such as *Heligmosomoides polygyrus bakeri* (*H. polygyrus*). Whether this is due to compensatory mechanisms is unclear. The transcription factor Bhlhe40 has been shown to control GM-CSF production and is proposed to be a novel regulator of T helper type 2 cells. We have found that Bhlhe40 is required in T cells for a protective memory response to secondary *H. polygyrus* infection. *H. polygyrus* rechallenge elicited dramatic Bhlhe40-dependent changes in gene and cytokine expression by lamina propria CD4^+^ T cells and *in vitro*-polarized T_H_2 cells, including induction of GM-CSF and maximal production of type 2 cytokines including IL-5. β_C_ chain-deficient, but not GM-CSF-deficient, mice rechallenged with *H. polygyrus* had severely impaired protective immunity. Our results demonstrate that Bhlhe40 is an essential regulator of T_H_2 cell immunity during helminth infection and reveal unexpected redundancy of β_C_ chain-dependent cytokines.

## INTRODUCTION

Helminthic worms are parasites of eukaryotic organisms which manipulate the immune system to establish chronic infections in diverse sites, often resulting in significant tissue damage (*1–4*). A stereotypical helminth infection results in production of alarmins including IL-25, IL-33, and thymic stromal lymphopoietin (TSLP), which in turn stimulate the development of type 2 immunity characterized by production of IL-4, IL-5, and IL-13 (*1, 2, 4*). The helminth *Heligmosomoides polygyrus bakeri* (hereafter *H. polygyrus*) is a natural pathogen of mice and has been frequently employed as a model helminth infection (*1–3, 5*). Several cytokines contribute to protective immunity against *H. polygyrus*, including IL-4 and IL-25, while IL-5, IL-33, and TSLP are considered dispensable for control of this infection (*6–9*).

However, in many instances of type 2 immunity, IL-5 plays a crucial role (*10, 11*), including in some helminth infections (*2, 4, 12*). Controversy previously existed as to whether IL-5 was protective against helminths, but it is now appreciated that deficiency in IL-5 only affects immunity to select parasites and sometimes in a life cycle stage-dependent manner (*4, 12*). IL-5 binds to the unique IL-5Rα chain paired with the common beta (β_C_) chain, which is shared with the receptors for GM-CSF and IL-3 (*13, 14*). This raises the possibility that redundancy within this cytokine family could render individual members dispensable during helminth infection. While IL-3 signaling is maintained in mice in the absence of the β_C_ chain (*Csf2rb*) through a unique beta chain (βIL3, *Csf2rb2*), GM-CSF and IL-5 signaling is abrogated in *Csf2rb*^-/-^ mice (*13, 14*). Despite absolute dependence on this shared receptor chain, which is largely responsible for downstream signaling (*13*), known roles for IL-5 and GM-CSF are divergent. GM-CSF has been proposed to be a central regulator of inflammation driven by T_H_1 and T_H_17 cells and affects both autoimmunity and infection (*15–23*). GM-CSF also contributes to pathological T_H_2 responses to allergens (*24–26*). However, little is known regarding GM-CSF during type 2 infections, though it is thought to be dispensable during infection with the helminth *Nippostrongylus brasiliensis* (*27*).

The transcription factor Bhlhe40 functions in B cells, NKT cells, T cells, and tissue-resident macrophages (*28–37*). We and others have described a crucial role for Bhlhe40 as a modulator of T_H_1 and T_H_17 cell cytokine production in infection and autoimmunity, particularly by controlling GM-CSF and IL-10 production (*32–36, 38*). Recent studies of *in vitro*-polarized T_H_2 cells (*39*) and house dust mite (HDM)-elicited airway T_H_2 cells (*40*) have identified Bhlhe40 as a potential regulator of this cell type *in vivo*. We have also recently described a role for Bhlhe40 in large peritoneal macrophages (LPMs) during type 2 immunity (*37*). Taken together, these data suggest that Bhlhe40 may regulate type 2 infections, possibly by controlling cytokine production from T cells or via a myeloid cell-intrinsic role.

Herein, we have found that Bhlhe40 and the β_C_ chain are essential to protective memory to secondary *H. polygyrus* infection. *Bhlhe40*^-/-^ mice exhibited severe defects during rechallenge infection with *H. polygyrus* and altered intestinal pathology, which were recapitulated in *Cd4-Cre*^+^ *Bhlhe40*^fl/fl^, but not *LysM-Cre*^+^ *Bhlhe40*^fl/fl^ mice. We defined a helminth-induced gene signature in lamina propria CD4^+^ T cells, which was disrupted in the absence of Bhlhe40. *In vitro*-polarized T_H_2 cells and *H. polygyrus*-elicited CD4^+^ T cells from the small intestine lamina propria exhibited Bhlhe40-dependent production of GM-CSF, and Bhlhe40-deficient CD4^+^ T cells also exhibited reduced production of IL-5 and other cytokines. Secondary infection of *Csf2rb*^-/-^, but not *Csf2*^-/-^, mice resulted in severely impaired protective immunity and altered intestinal pathology. Overall, Bhlhe40 serves as a pivotal regulator of the T_H_2 cell transcriptional response to helminth infection, in part by modulating GM-CSF and IL-5 production, and reveals redundant roles for these cytokines not observed during deficiency of either factor alone.

## RESULTS

### Bhlhe40 is required for control of *H. polygyrus* rechallenge

Primary oral challenge of C57BL/6 mice with *H. polygyrus* results in chronic infection, but rechallenge evokes a protective recall response which limits infection (*41*). To assess whether Bhlhe40 was required for immunity to *H. polygyrus*, we challenged C57BL/6 *Bhlhe40*^+/+^ and *Bhlhe40*^-/-^ mice with infective *H. polygyrus* larvae (L3), cured them by treatment with pyrantel pamoate, and rechallenged them with infective L3. While *Bhlhe40*^+/+^ and *Bhlhe40*^-/-^ mice exhibited similar parasite fecal egg burdens during primary infection, *Bhlhe40*^-/-^ mice had a much higher egg burden during secondary infection as compared to *Bhlhe40*^+/+^ mice (Fig. 1A). While there was a trend towards increased adult worm burden in *Bhlhe40*^-/-^ as compared to *Bhlhe40*^+/+^ mice after rechallenge (Fig. 1B), this did not reach statistical significance, indicating that their increased egg burden was primarily due to increased worm fecundity. *H. polygyrus*-rechallenged *Bhlhe40*^+/+^ and *Bhlhe40*^-/-^ mice had dramatically different pathology. Type 2 granulomas can form around developing parasites and have been correlated with protective immunity (*41*). Small intestines from *Bhlhe40*^+/+^ mice exhibited many granulomas, while those from *Bhlhe40*^-/-^ mice resembled healthy tissue (Fig. 1, C and D). Histological analysis and immunostaining showed reduced immune infiltration and damage to the smooth muscle layer in *H. polygyrus*-rechallenged *Bhlhe40*^-/-^ as compared to *Bhlhe40*^+/+^ mice (Fig. 1, E and F).

**Figure 1.**
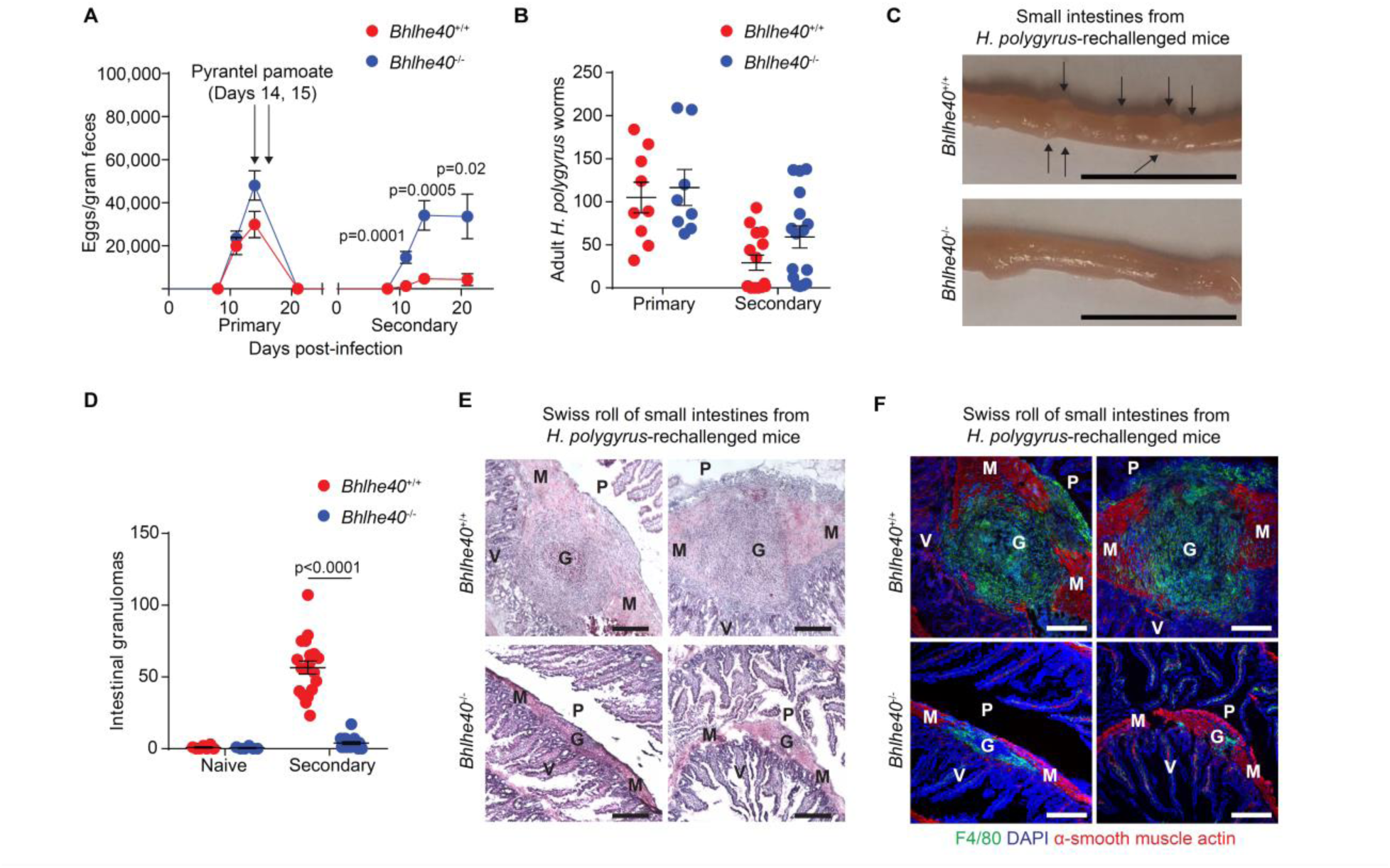
Bhlhe40 is required for a protective recall response to *H. polygyrus*. **(A-C)** *H. polygyrus-*infected *Bhlhe40*^+/+^ and *Bhlhe40*^-/-^ mice were analyzed for **(A)** quantitation of *H. polygyrus* eggs/gram feces over time during secondary infection, **(B)** quantitation of adult worms recovered from the intestines of mice experiencing primary or secondary infection, and **(C)** small intestine morphology after secondary infection. Arrows point to granulomas. Scale bar, 1 cm. **(D)** Quantitation of intestinal granulomas. **(E and F)** *H. polygyrus-*rechallenged *Bhlhe40*^+/+^ and *Bhlhe40*^-/-^ mice were analyzed histologically on adjacent sections by **(E)** hematoxylin and eosin staining and **(F)** immunostaining for F4/80, α-SMA, and DAPI performed on swiss rolls of the proximal small intestine (2 representative lesions from each genotype). G, granuloma. M, muscle layers. P, peritoneal space. V, villi. Scale bar, 200 μm. Data are representative of at least 3 independent experiments **(C, E, F)** or are pooled from at least 3 independent experiments **(A, B, D)**. Data are mean ± s.e.m. Significance calculated with an unpaired Student’s *t*-test.

When we explored the cellular composition of the small intestine lamina propria (SILP) following *H. polygyrus* rechallenge of *Bhlhe40*^+/+^ mice by flow cytometry, we found both CD45^+^CD64^+^F4/80^+^MHC-II^+^Ly6C^-^ resident macrophages (*42, 43*) and another CD45^+^F4/80^+^CD64^+^MHC-II^lo^Ly6C^lo^ autofluorescent population, which we termed granuloma-associated monocytes/macrophages (GMMs) (Fig. 2, A and B). This latter population may correspond to previously described clodronate-sensitive alternatively activated macrophages seen by immunostaining during *H. polygyrus* infection (*44*). SILP CD45^+^F4/80^+^CD64^-^CD11b^+^SSC-A^hi^ eosinophils were also significantly increased after secondary *H. polygyrus* infection (Fig. 2, A and B and fig. S1A). However, GMMs and eosinophils were greatly reduced in *H. polygyrus*-rechallenged *Bhlhe40*^-/-^ as compared to *Bhlhe40*^+/+^ mice, in contrast to SILP CD3^+^ T cells (Fig. 2, A and B and fig. S1B). Because *H. polygyrus* infection is known to change the cellular composition of the peritoneal cavity (*4, 45, 46*), we assessed accumulation of LPMs and peritoneal eosinophils, and found severe reductions in both populations in *H. polygyrus*-rechallenged *Bhlhe40*^-/-^ as compared to *Bhlhe40*^+/+^ mice, as well as impaired LPM polarization as assessed by RELMα staining (Fig. 2, C to E and fig. S1, C to E). In contrast, serum *H. polygyrus*-specific IgG1 titers were not affected by loss of Bhlhe40 (fig. S1F). These data indicated impaired responses by multiple myeloid cell lineages to *H. polygyrus* rechallenge in *Bhlhe40*^-/-^ mice.

**Figure 2.**
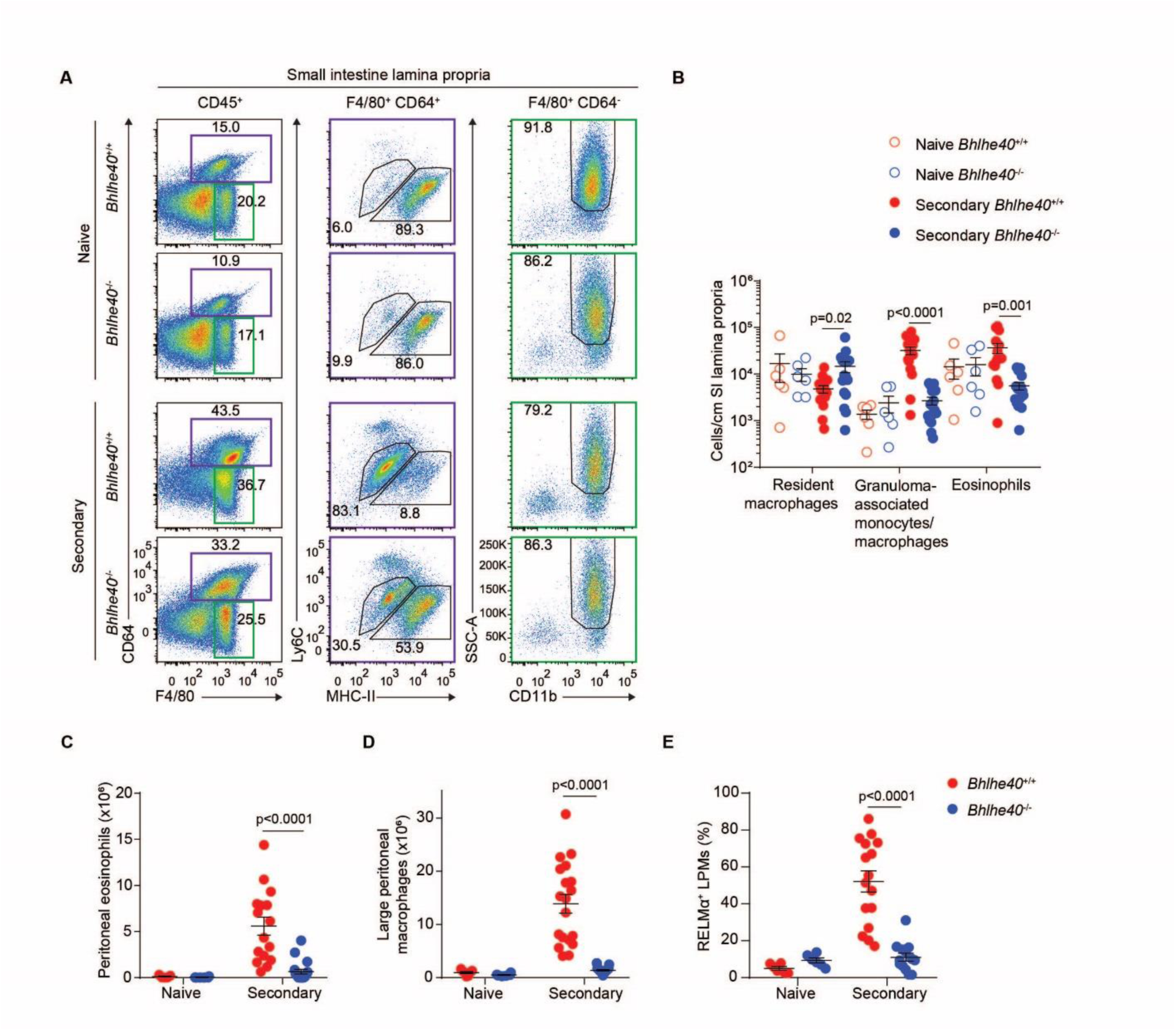
Bhlhe40 is required for normal myeloid cell responses to *H. polygyrus* rechallenge. **(A and B)** Naïve and *H. polygyrus-*rechallenged *Bhlhe40*^+/+^ and *Bhlhe40*^-/-^ mice were analyzed by flow cytometry for **(A)** SILP myeloid cells and **(B)** quantitation as in **(A)**. **(C-E)** Naïve and *H. polygyrus-*rechallenged *Bhlhe40*^+/+^ and *Bhlhe40*^-/-^ mice were analyzed by flow cytometry for quantitation of **(C)** peritoneal eosinophils, **(D)** LPMs, and **(E)** the frequency of RELMα^+^ LPMs. Data are representative of at least 3 independent experiments **(A)** or are pooled from at least 3 independent experiments **(B-E)**. Data are mean ± s.e.m. Significance calculated with an unpaired Student’s *t*-test.

### Bhlhe40 is required in T cells for a normal immune response to *H. polygyrus*

We next employed *Cd4-Cre*^+^ *Bhlhe40*^fl/fl^ and *LysM-Cre*^+^ *Bhlhe40*^fl/fl^ mice to address whether loss of Bhlhe40 specifically in T cells or myeloid cells recapitulated the phenotype of *Bhlhe40*^-/-^ mice during secondary *H. polygyrus* infection. *Cd4-Cre*^+^ *Bhlhe40*^fl/fl^ mice were severely impaired in controlling *H. polygyrus* rechallenge and lacked intestinal granulomas as compared to *Cd4-Cre*^-^ *Bhlhe40*^fl/fl^ mice (Fig. 3, A and B). When we assessed the response of myeloid cells to secondary infection, we found that loss of Bhlhe40 selectively in T cells was sufficient to perturb both the SILP and peritoneal responses (Fig. 3, C to F). In contrast, *LysM-Cre*^+^ *Bhlhe40*^fl/fl^ mice were able to control infection and formed intestinal granulomas comparably to *LysM-Cre*^-^ *Bhlhe40*^fl/fl^ mice (Fig. 3, G and H). Therefore, Bhlhe40 expression was required in T cells to control secondary *H. polygyrus* infection, but was dispensable in LysM-expressing myeloid cells.

**Figure 3.**
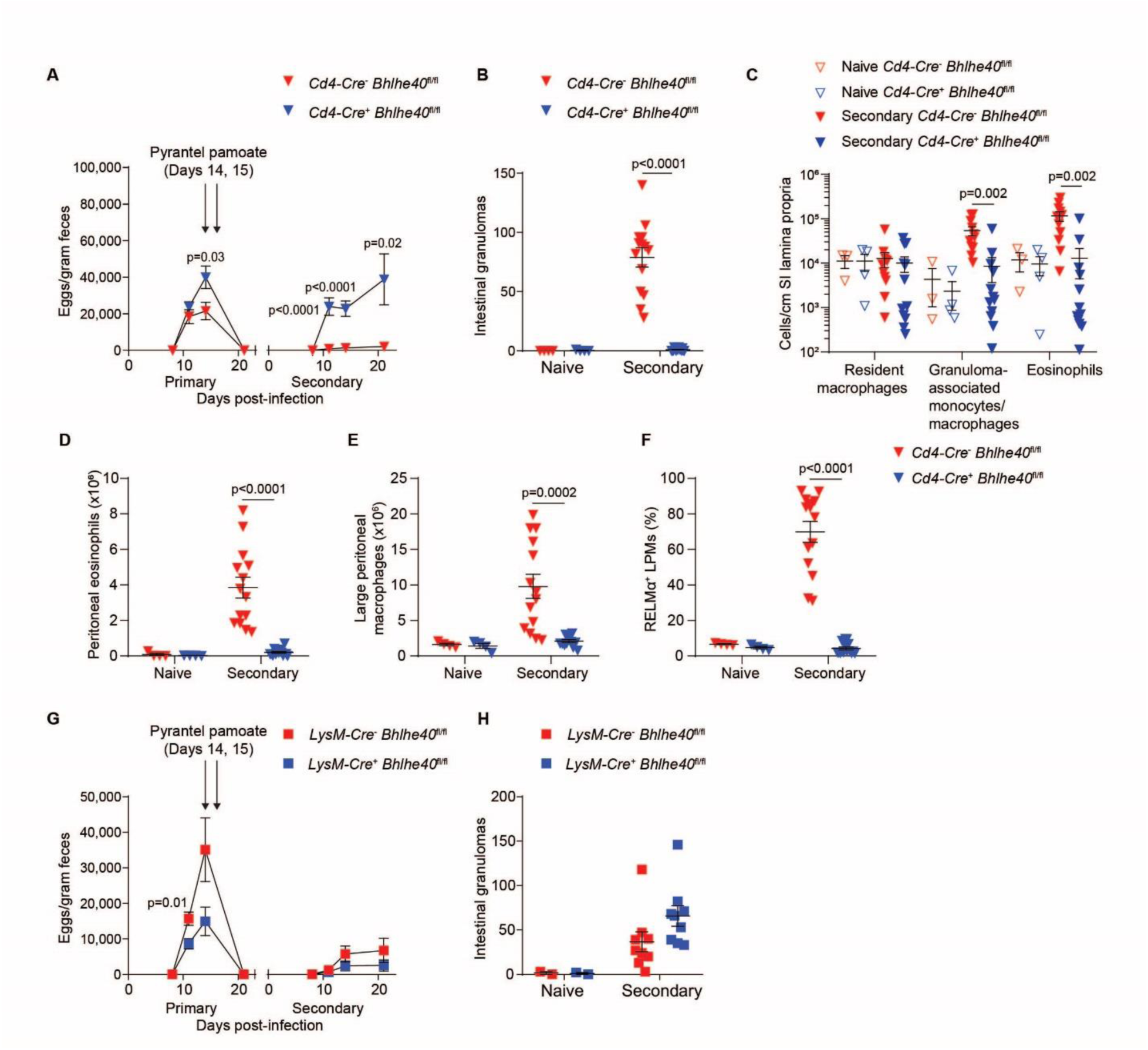
Bhlhe40 is required in T cells for a normal memory response to *H. polygyrus*. **(A)** *H. polygyrus-*rechallenged *Cd4-Cre*^-^ *Bhlhe40*^fl/fl^ and *Cd4-Cre*^+^ *Bhlhe40*^fl/fl^ mice were analyzed for quantitation of *H. polygyrus* eggs/gram feces over time. **(B)** Naïve and *H. polygyrus-* rechallenged *Cd4-Cre*^-^ *Bhlhe40*^fl/fl^ and *Cd4-Cre*^+^ *Bhlhe40*^fl/fl^ mice were analyzed for quantitation of intestinal granulomas. **(C-F)** Naïve and *H. polygyrus-*rechallenged *Cd4-Cre*^-^ *Bhlhe40*^fl/fl^ and *Cd4-Cre*^+^ *Bhlhe40*^fl/fl^ mice were analyzed by flow cytometry for quantitation of **(C)** SILP myeloid cells, **(D)** peritoneal eosinophils, **(E)** LPMs, and **(F)** the frequency of RELMα^+^ LPMs. **(G)** *H. polygyrus-*rechallenged *LysM-Cre*^-^ *Bhlhe40*^fl/fl^ and *LysM-Cre*^+^ *Bhlhe40*^fl/fl^ mice were analyzed for quantitation of *H. polygyrus* eggs/gram feces over time. **(H)** Naïve and *H. polygyrus-*rechallenged *LysM-Cre*^-^ *Bhlhe40*^fl/fl^ and *LysM-Cre*^+^ *Bhlhe40*^fl/fl^ mice were analyzed for quantitation of intestinal granulomas. Data are pooled from 3 independent experiments **(A-F)** or 2 independent experiments **(G and H)**. Data are mean ± s.e.m. Significance calculated with an unpaired Student’s *t*-test.

### Bhlhe40 is required for a normal CD4^+^ T cell transcriptional response to *H. polygyrus*

We next asked whether loss of Bhlhe40 dysregulated CD4^+^ T cell gene expression in response to *H. polygyrus* rechallenge. To address this, we sorted CD4^+^ T cells from the SILP of naïve and rechallenged *Bhlhe40*^+/+^ and *Bhlhe40*^-/-^ mice for gene expression microarrays. By comparing CD4^+^ T cells from naïve and *H. polygyrus*-rechallenged *Bhlhe40*^+/+^ mice, we defined a helminth-induced signature which included transcripts for cytokines (including *Areg*, *Il3*, *Il4*, *Il5*, *Il6*, *Il13*, *Csf1*, *Csf2, Lif*, *Tnf*, *Tnfsf11*), cytokine receptors (including *Il1rl1*, *Il1r2*, *Il17rb*), and transcription factors (including *Atf3*, *Bhlhe40*, *Gata3*, *Nfil3*, *Pparg*, *Rbpj*, *Vdr*, and *Zeb2*) (Fig. 4A and fig. S2A). When we assessed Bhlhe40-dependent genes after *H. polygyrus* rechallenge, we found that a significant majority were part of the helminth-induced signature and that Bhlhe40-dependent genes were distinct in CD4^+^ T cells from naïve and *H. polygyrus*-rechallenged mice (Fig. 4, B and C and fig. S2, B and C). When we used gene set enrichment analysis (GSEA) to look for Bhlhe40-dependent gene modules, we noted that two of the most enriched sets in *Bhlhe40*^+/+^ as compared to *Bhlhe40*^-/-^ CD4^+^ T cells after *H. polygyrus* rechallenge were “growth factor activity” and “cytokine activity,” reflecting altered expression of cytokine genes including *Areg*, *Il5*, *Il6*, *Il13*, *Csf1*, *Csf2*, and *Lif*, but not *Il3*, *Il4*, or *Tnf* (Fig. 4, D and E). Furthermore, when we assessed differential expression of the lineage-specifying transcription factors of each T helper cell subset as well as a recently defined set of transcriptional regulators of *in vitro*-polarized T_H_2 cells (*39*), we observed reduced expression of *Pparg* in *Bhlhe40*^-/-^ as compared to *Bhlhe40*^+/+^ CD4^+^ T cells from *H. polygyrus*-rechallenged mice (2.4-fold reduced), but only subtle changes in other factors including *Gata3* (1.4-fold reduced) (Fig. 4E). *Bhlhe40* expression was markedly induced in mice experiencing secondary infection as compared to naïve mice (Fig. 4E). Using *Bhlhe40*^GFP^ bacterial artificial chromosome reporter mice (*34*), we observed a marked increase in SILP GFP^+^ CD4^+^ T cells after *H. polygyrus* rechallenge (Fig. 4F). Taken together, these data showed that Bhlhe40 was induced by SILP CD4^+^ T cells in response to *H. polygyrus* rechallenge and was critical for their normal transcriptional program.

**Figure 4.**
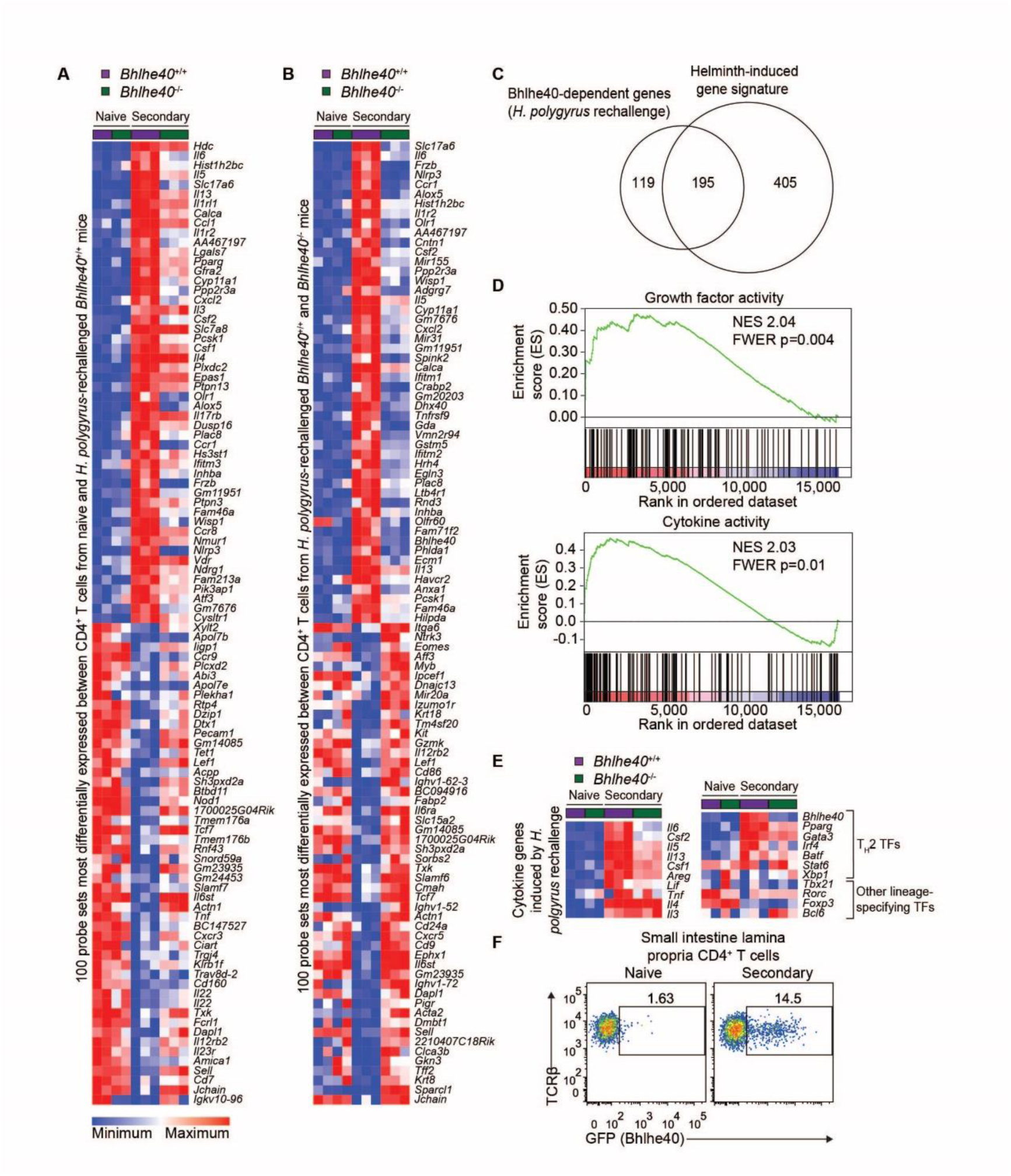
Loss of Bhlhe40 dysregulates the CD4^+^ T cell transcriptional response to *H. polygyrus* rechallenge. **(A and B)** Gene expression microarray data were analyzed for the 100 most differentially expressed probe sets in SILP CD4^+^ T cells from **(A)** naïve and *H. polygyrus*-rechallenged *Bhlhe40*^+/+^ mice and **(B)** *H. polygyrus*-rechallenged *Bhlhe40*^+/+^ and *Bhlhe40*^-/-^ mice. **(C)** Gene expression microarray data were analyzed for shared and unique Bhlhe40-dependent genes (≥2-fold differentially expressed between SILP CD4^+^ T cells from *H. polygyrus*-rechallenged *Bhlhe40*^+/+^ and *Bhlhe40*^-/-^ mice) with the helminth-induced signature (≥2-fold differentially expressed between SILP CD4^+^ T cells from naïve and *H. polygyrus*-rechallenged *Bhlhe40*^+/+^ mice), depicted as a Venn diagram. **(D)** GSEA of gene expression microarray data for selected C5 gene sets enriched in *Bhlhe40*^+/+^ versus *Bhlhe40*^-/-^ SILP CD4^+^ T cells from *H. polygyrus*-rechallenged mice. NES, normalized enrichment score. FWER, family-wise error rate. **(E)** Gene expression microarray data were analyzed for (left) expression of cytokines induced by *H. polygyrus* rechallenge and (right) expression of T_H_2 and T helper cell lineage-specifying transcription factors in SILP CD4^+^ T cells from naïve and *H. polygyrus*-rechallenged *Bhlhe40*^+/+^ and *Bhlhe40*^-/-^ mice. TF, transcription factor. **(F)** Flow cytometry of *Bhlhe40*^GFP^ transgene reporter expression in SILP CD4^+^ T cells from *Bhlhe40*^GFP+^ mice. Data in **(F)** are representative of 2 independent experiments. Microarray data are from a single experiment.

### Loss of Bhlhe40 impairs T cell cytokine production in response to *H. polygyrus*

Next, we restimulated SILP cells from naïve and rechallenged *Cd4-Cre*^-^ *Bhlhe40*^fl/fl^ and *Cd4-Cre*^+^ *Bhlhe40*^fl/fl^ mice *ex vivo* with phorbol 12-myristate 13-acetate (PMA) and ionomycin to assess cytokine production. Rechallenge with *H. polygyrus* induced production of IL-4, IL-5, and IL-13 from *Cd4-Cre*^-^ *Bhlhe40*^fl/fl^ and *Cd4-Cre*^+^ *Bhlhe40*^fl/fl^ CD4^+^ T cells; however, Bhlhe40 was required for normal frequencies of single- and multi-cytokine-producing cells, largely due to reductions in IL-5^+^ and IL-13^+^ CD4^+^ T cells (Fig. 5, A to C). Because *Csf2* was upregulated by *H. polygyrus* rechallenge in a Bhlhe40-dependent fashion, we also assessed GM-CSF and found that it was markedly induced by rechallenge (∼35% of *Cd4-Cre*^-^ *Bhlhe40*^fl/fl^ CD4^+^ T cells) and that this required Bhlhe40 (∼10% of *Cd4-Cre*^+^ *Bhlhe40*^fl/fl^ CD4^+^ T cells) (Fig. 5, D and E). T_H_2 cells disseminate widely in mice infected with *H. polygyrus* (*45, 46*). CD4^+^ T cell cytokine responses in the peritoneal cavity and mesenteric lymph nodes from *H. polygyrus*-rechallenged *Cd4-Cre*^-^ *Bhlhe40*^fl/fl^ and *Cd4-Cre*^+^ *Bhlhe40*^fl/fl^ mice were generally consistent with those in the SILP (fig. S3). To assess whether Bhlhe40 was also required for normal cytokine production by a pure population of T_H_2 cells, we differentiated naïve splenic CD4^+^ T cells into T_H_2 cells and restimulated them to assess cytokine production. We found that Bhlhe40 was essential for production of GM-CSF and that loss of Bhlhe40 also impaired *in vitro* production of type 2 cytokines (fig. S4). Therefore, these data indicated that Bhlhe40 is required *in vitro* and *in vivo* for normal T_H_2 cell function.

**Figure 5.**
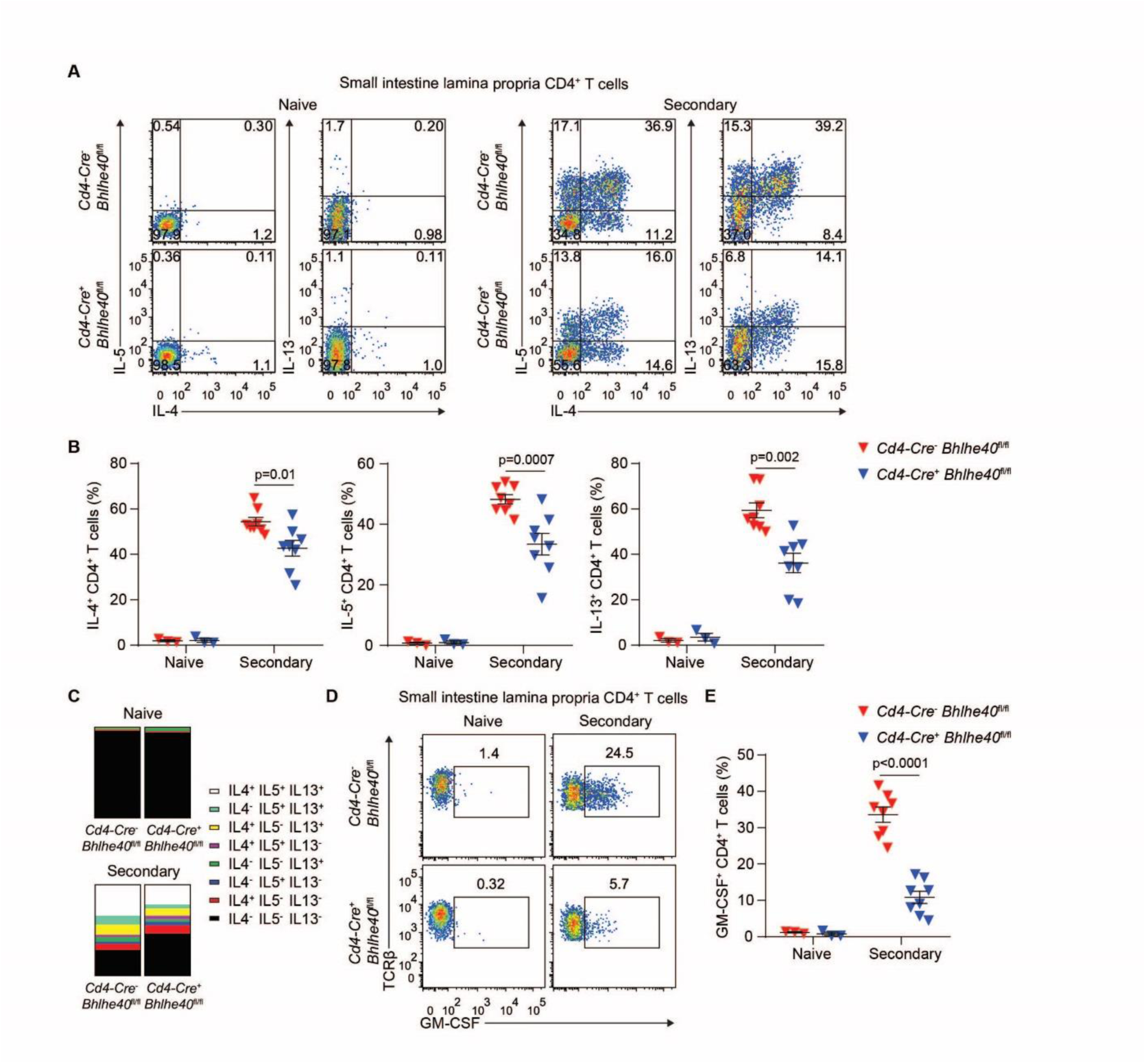
Bhlhe40 is required for normal CD4^+^ T cell production of β_C_ chain-dependent cytokines. **(A-C)** Naïve and *H. polygyrus-*rechallenged *Cd4-Cre*^-^ *Bhlhe40*^fl/fl^ and *Cd4-Cre*^+^ *Bhlhe40*^fl/fl^ mice were analyzed by flow cytometry for **(A)** IL-4-, IL-5-, and IL-13-producing CD4^+^ T cells, **(B)** quantitation of the frequency of IL-4^+^, IL-5^+^, and IL-13^+^ CD4^+^ T cells, and **(C)** quantitation of the frequency of CD4^+^ T cells producing one or more cytokines after *in vitro* PMA/ionomycin stimulation of SILP cells. **(D and E)** Naïve and *H. polygyrus-* rechallenged *Cd4-Cre*^-^ *Bhlhe40*^fl/fl^ and *Cd4-Cre*^+^ *Bhlhe40*^fl/fl^ mice were analyzed by flow cytometry for **(D)** GM-CSF-producing CD4^+^ T cells and **(E)** quantitation as in **(D)** after *in vitro* PMA/ionomycin stimulation of SILP cells. Data are representative of 2 independent experiments **(A and D)** or are pooled from 2 independent experiments **(B, C, E)**. Data are mean ± s.e.m. Significance calculated with an unpaired Student’s *t*-test.

As Bhlhe40 is a known repressor of IL-10 (*32–34, 36*), we also assessed IL-10 production from SILP CD4^+^ T cells. Indeed, SILP CD4^+^ T cells lacking Bhlhe40 produced significantly higher levels of IL-10 after *H. polygyrus* rechallenge (fig. S5A). Nevertheless, genetic deletion of IL-10 in *Bhlhe40*^-/-^ *Il10*^-/-^ mice was not sufficient to restore control of infection or normal intestinal granulomatous pathology (fig. S5, B and C). Collectively, these data demonstrated a key role for Bhlhe40 in CD4^+^ T cell cytokine responses to helminth infection, notably controlling production of the β_C_ chain family cytokines IL-5 and GM-CSF.

### Loss of the β_C_ chain impairs protective memory to *H. polygyrus*

As protective memory responses to *H. polygyrus* rechallenge are unaffected by IL-5 blockade (*6*), we first asked whether loss of GM-CSF signaling was sufficient to impair control of a secondary infection with *H. polygyrus*. Genetic deletion of GM-CSF (*Csf2*^-/-^ mice) did not result in severe defects in control of *H. polygyrus* infection as compared to *Csf2*^+/+^ mice (fig. S6, A to F). As IL-5 and GM-CSF were individually dispensable, we then rechallenged *Csf2rb*^+/+^ and *Csf2rb*^-/-^ mice. Remarkably, *Csf2rb*^-/-^ mice had a severe defect in control of *H. polygyrus* as compared to *Csf2rb*^+/+^ mice (Fig. 6A). *Csf2rb*^-/-^ mice did not form intestinal granulomas or develop a normal SILP myeloid cell response to *H. polygyrus* rechallenge as compared to *Csf2rb*^+/+^ mice (Fig. 6, B and C). Furthermore, loss of the β_C_ chain resulted in defects in peritoneal myeloid cell responses similar to those seen in *Bhlhe40*^-/-^ and *Cd4-Cre*^+^ *Bhlhe40*^fl/fl^ mice (Fig. 6, D to F). To exclude a role for β_C_ chain-dependent IL-3 signaling, we singly blocked GM-CSF or IL-5 signaling with neutralizing antibodies or blocked both cytokines together. Double, but not single, blockade resulted in severely impaired protective immunity (fig. S6, G and H). Taken together, these data indicated that IL-5 and GM-CSF were collectively, but not individually, critical to control of *H. polygyrus* rechallenge.

**Figure 6.**
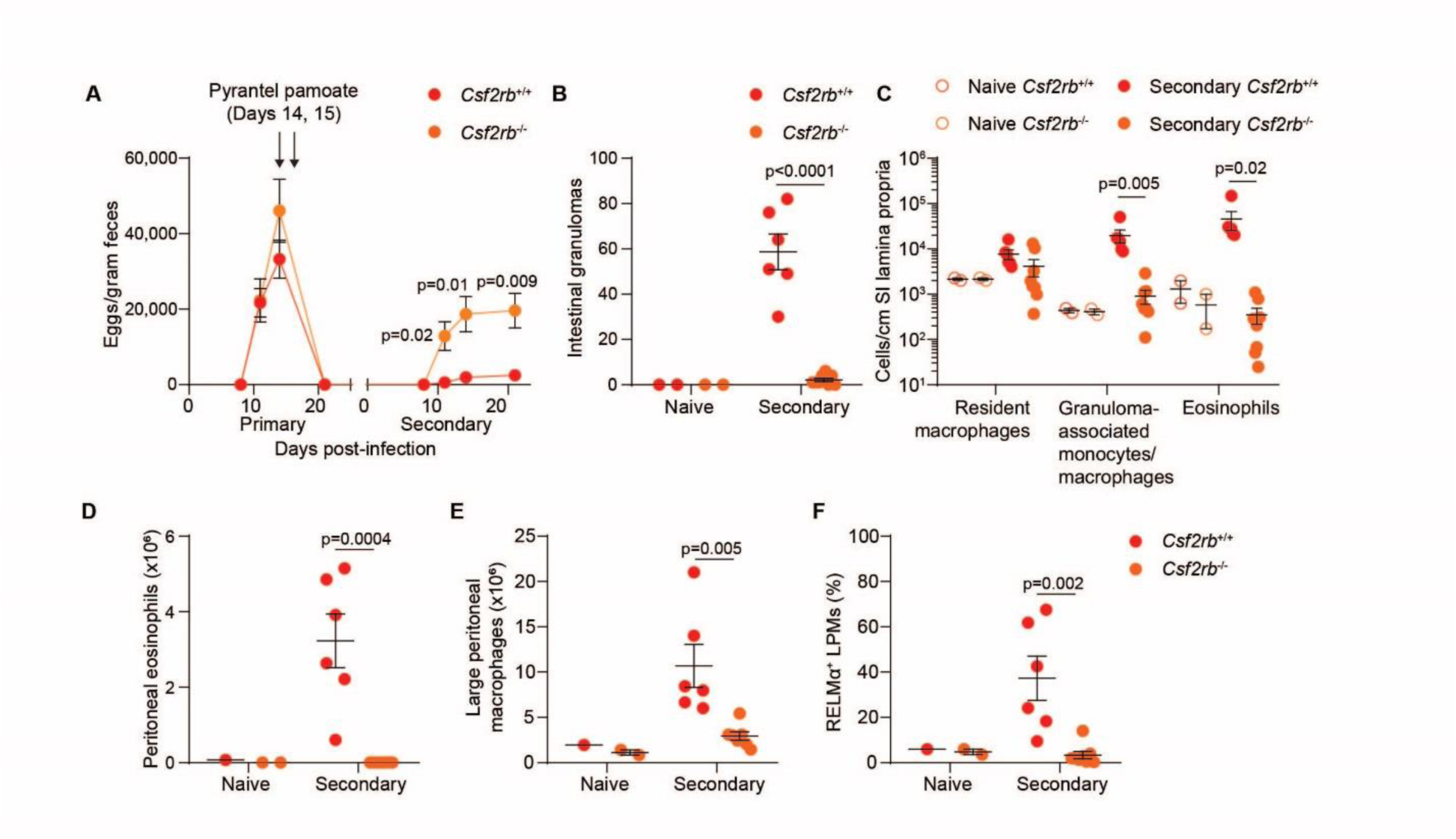
The β_C_ chain is necessary for control of *H. polygyrus* rechallenge. **(A)** *H. polygyrus-*rechallenged *Csf2rb*^+/+^ and *Csf2rb*^-/-^ mice were analyzed for quantitation of *H. polygyrus* eggs/gram feces over time. **(B)** Naïve and *H. polygyrus-*rechallenged *Csf2rb*^+/+^ and *Csf2rb*^-/-^ mice were analyzed for quantitation of intestinal granulomas. **(C-F)** Naïve and *H. polygyrus-*rechallenged *Csf2rb*^+/+^ and *Csf2rb*^-/-^ mice were analyzed by flow cytometry for quantitation of **(C)** SILP myeloid cells, **(D)** peritoneal eosinophils, **(E)** LPMs, and **(F)** the frequency of RELMα^+^ LPMs. Data are pooled from 2 independent experiments. Data are mean ± s.e.m. Significance calculated with an unpaired Student’s *t*-test.

## DISCUSSION

We have demonstrated that protective memory responses against *H. polygyrus* are critically dependent on Bhlhe40 and the β_C_ chain. Bhlhe40 is specifically required in CD4^+^ T cells to promote a normal myeloid cell response to *H. polygyrus* by supporting expression of cytokine transcripts as well as other potential regulators of helminth infection, including *Areg*, *Csf2*, *Il5*, *Il13*, *Nlrp3*, and *Pparg* (*47–49*). We have also described the CD4^+^ T cell transcriptome within the SILP during secondary *H. polygyrus* infection. These data provide unique insight into the CD4^+^ T cell global transcriptional response to helminths at the site of infection. We found that Bhlhe40 is a key regulator of T_H_2 cell cytokine production. Notably, GM-CSF and IL-5 were markedly stimulated by *H. polygyrus* rechallenge and this response was Bhlhe40-dependent. In light of these data, we assessed the importance of the β_C_ chain during *H. polygyrus* rechallenge and found that it was critically required for protective immunity, despite control of secondary *H. polygyrus* infection during IL-5 blockade (this manuscript and (*6*)), during GM-CSF blockade, and in *Csf2*^-/-^ mice. These data support redundant roles for Bhlhe40-dependent β_C_ chain-dependent cytokines in protective immunity to *H. polygyrus*.

When we compared differentially expressed transcripts between *H. polygyrus*-elicited and naive SILP CD4^+^ T cells, we found a remarkable similarity to the transcriptional profile of HDM-elicited airway T_H_2 cells (*Bhlhe40*, *Cd200r1*, *Il6*, *Plac8*, *Igfbp7*) (*40*), demonstrating significant conservation of the T_H_2 transcriptional program independent of tissue environment and stimulus. When we assessed whether Bhlhe40 was functionally required in T cells during type 2 immunity, we found that SILP CD4^+^ T cells activated by *H. polygyrus* rechallenge required Bhlhe40 for normal expression of many helminth-induced genes and to control helminth rechallenge. We therefore demonstrate for the first time that Bhlhe40 regulates *in vivo* T_H_2 cell responses, consistent with a recent screen for novel regulators of *in vitro*-polarized T_H_2 cells that identified Bhlhe40 (*39*). Notably, *Pparg* expression was reduced in the absence of Bhlhe40, and as PPARγ is required for normal T_H_2 cell responses and protective immunity to *H. polygyrus*, this may indicate that some of the effects of Bhlhe40 deficiency are indirect (*47, 50*). Our work and that of others has now established key roles for Bhlhe40 in TH1, TH2, and T_H_17 cells as a pivotal regulator of GM-CSF, IL-10, and other cytokines (*32–36, 38*). These data and a recent study on c-Maf (*51*) suggest that T cell production of many cytokines may be controlled by multi-lineage transcription factors in addition to lineage-restricted factors. In light of the critical role for Bhlhe40 in multiple CD4^+^ T cell subsets, clinical targeting of factors regulating Bhlhe40 may possess significant therapeutic potential.

While GM-CSF and IL-5 have long been known to share common signaling through the β_C_ chain, their described functions are largely distinct, with GM-CSF contributing to T_H_1 and T_H_17 cell-driven inflammation and IL-5 contributing to type 2 immunity (*10, 11, 16, 52*). We have demonstrated that protective memory to a helminth infection unaffected by IL-5 blockade and also insensitive to GM-CSF deficiency is nonetheless dependent on the combination of GM-CSF and IL-5 signaling through the β_C_ chain. It remains to be determined how GM-CSF and IL-5 compensate for each other, whether by direct substitution or via effects on distinct arms of the type 2 response. While redundancy between β_C_ chain family cytokines is not well described, it is known that these cytokines can regulate eosinophils in a complementary manner (*53*). Future studies should establish whether GM-CSF and IL-5 are collectively involved in protective immunity to other helminth infections. As β_IL3_ chain-dependent IL-3 signaling is preserved in *Csf2rb*^-/-^ mice (*13, 14*), it is also of interest to establish whether additionally blocking IL-3 results in a more severe defect in immunity to *H. polygyrus* than is observed in *Csf2rb*^-/-^ mice. IL-3 regulates basophilia in response to *H. polygyrus* and basophils help control infection with this helminth (*54, 55*), suggesting that IL-3 is likely important for control of *H. polygyrus* infection.

Our findings reveal novel transcriptional and cytokine regulation of the immune response to helminth infection, elucidating a critical function for Bhlhe40 during *in vivo* T_H_2 cell immunity and an unexpected role for β_C_ chain-dependent signaling. Our data suggest that the importance of β_C_ chain family cytokines may have been underestimated in type 2 diseases because of redundancy. Further studies are needed to elucidate whether compensation between β_C_ chain family cytokines is a general theme of type 2 immunity and whether combinatorial targeting of these factors could yield improved clinical outcomes in diseases of pathological type 2 immunity.

## MATERIALS AND METHODS

### Mice

C57BL/6 (Taconic and Jackson), *Il10*^-/-^ (B6.129P2-Il10tm1Cgn/J, Jackson), *Csf2*^-/-^ (B6.129S-*Csf2^tm1Mlg^*/J, Jackson), *Csf2rb*^-/-^ (B6.129S1-*Csf2rb^tm1Cgb^*/J, Jackson), *Cd4-Cre* (B6.Cg-Tg(Cd4-cre)1Cwi/BfluJ, Jackson) and *LysM-Cre* (B6N.129P2(B6)-*Lyz2^tm1(cre)Ifo^*/J, Jackson) mice were obtained from the vendors listed. *Bhlhe40*^-/-^ (10 generations backcrossed to the C57BL/6 background) (*33, 56*), *Bhlhe40*^GFP+^ (10 generations backcrossed to the C57BL/6 background) (*34*), and *Bhlhe40*^fl/fl^ (*32*) mice have been previously reported. All mice were maintained in our specific pathogen free animal facility. Sex-matched littermates were used for experiments whenever possible. Experiments with *Csf2*^-/-^ and *Csf2rb*^-/-^ mice used cohoused, age-matched, sex-matched C57BL/6 controls bred in our animal facility or purchased from Jackson. All animal experiments were approved by the Animal Studies Committee of Washington University in St. Louis.

### Heligmosomoides polygyrus bakeri infections

Infective *Heligmosomoides polygyrus bakeri* third-stage larvae (L3) were prepared as described (*5*). For both primary and secondary infection, mice were orally gavaged with 200 L3 with a 20-gauge ball-tipped gavage needle. Female mice were used for experiments whenever possible. Fecal samples were collected at 8, 11, 14, and 21 days after primary and secondary infection and weighed. Eggs were counted by dissolution of feces in 5 ml water, followed by a 1:1 dilution with a saturated sodium chloride solution before loading into a McMaster counting chamber (Chalex LLC). For rechallenge experiments, mice were cleared of infection by oral gavage with 2 mg of pyrantel pamoate (Columbia Laboratories) diluted in Dulbecco’s PBS on days 14 and 15 after primary infection and were rested for 3-4 weeks before reinfection. Blood was collected at day 21 post-rechallenge and allowed to clot for >1 hour before serum collection by centrifugation at 1,500 g for 15 minutes. Mice were sacrificed on day 17 or 22 of secondary infection for assessment of cellular responses. Macroscopic granulomas present at day 22 of secondary infection that were visible to the naked eye were counted along the length of the intact intestines. Assessment of the intestinal worm burden was performed at day 17 or 22 of primary or secondary infection. Adult worms were collected by cutting the intestine open and into ∼4 in sections, placing the tissue and contents into a metal filter on top of a 50 ml tube of PBS, and setting the tube in a 37°C water bath. Worms then actively migrated through the filter over several hours and sedimented. Worms were counted visually in petri dishes.

### *In vivo* antibody blockade

Mice were infected, cleared, and rested as above. Immediately before reinfection, mice were injected i.p. with 300 µg of αGM-CSF (Leinco, G670, clone MP1-22E9), 300 ug of αIL-5 (BioXCell, BE0198, clone TRFK5), or control polyclonal rat IgG (Sigma Aldrich, I4131) and were given the same dose i.p. every other day until sacrifice (*52*).

### *In vitro* T_H_2 cell culture

The EasySep mouse naïve CD4+ T cell isolation kit (Stemcell Technologies, 19765A) was used to isolate T cells from the spleens of untreated mice. Cells were cultured in complete Iscove’s modified Dulbecco’s media (cIMDM, with added 10 % FBS, L-glutamine, sodium pyruvate, non-essential amino acids, penicillin/streptomycin, and β-mercaptoethanol) in flat-bottom, tissue culture-treated 48 well plates with plate-bound anti-CD3 and anti-CD28. Plates were coated overnight with anti-CD3 antibody (BioXCell, 2 μg/mL, clone 145-2C11) and anti-CD28 antibody (BioLegend, 2 μg/ml, clone 37.51) before use. To polarize naïve T cells to T_H_2 cells, IL-4 (BioLegend, 10 ng/mL) and anti-IFN-γ (Leinco, 5 μg/mL, clone H22) were added at the start of culture. Cells were split at day 3, and on day 4 cells were stimulated for 24 hours with plate-bound anti-CD3 and anti-CD28 (coated as above) for assessment of secreted cytokines in the supernatant by ELISA.

### Enzyme-linked immunosorbent assay (ELISA)

Nunc Maxisorp plates were coated with capture antibodies (for cytokine ELISAs) or *H. polygyrus* lysate (for *H. polygyrus*-specific IgG1 ELISA) overnight. Wells were washed with PBS-Tween 20 (0.05%), and 1% BSA in PBS was added for one hour to block. After washing, culture supernatants (cytokine ELISAs) or serum (IgG1 ELISA) were added for 1-2 hours, except for the GM-CSF ELISA (overnight). After washing, cytokine-detection biotinylated antibodies or horseradish peroxidase (HRP)-conjugated anti-mouse IgG1 antibody were added for 1 hour, except for the GM-CSF ELISA (2 hours). For cytokine ELISAs, following another wash, streptavidin-conjugated horseradish peroxidase (HRP) was added for 1 hour. After washing, a 1:1 mixture of room temperature TMB A and B solutions (BD OptEIA) was added. The reaction was stopped with 1 M H_3_PO_4_. Samples were analyzed on an iMark microplate reader (BioRad). For cytokine ELISAs, standard curves were generated with purified cytokines. For the IgG1 ELISA, serum was diluted between 10^-2^ and 10^-8^ and the last well with an OD_450_ above 0.100 represented the titer. ELISA reagents are listed in Table S1.

To generate *H. polygyrus* lysate, adult worms collected as above were washed repeatedly in PBS, and ground in a Dounce homogenizer in 1 mL of PBS. Debris was then pelleted by centrifugation at 16,000 g for 20 minutes at 4°C. The supernatant was passed through a 0.2 μm filter and stored at -80°C. Lysate was titrated for an ELISA with the serum of *H. polygyrus*-rechallenged mice, and the greatest dilution of lysate (1:1000) which yielded maximal signal was used.

### Brightfield and epifluorescence microscopy

The proximal 6 cm of the small intestine were formed into a swiss roll (*57*) and were frozen in OCT media. 10 μm cryosections were cut using a Leica CM1950 cryostat. For hematoxylin and eosin staining, sections were fixed in 4% PFA, washed, and stained. For immunofluorescent staining, sections were fixed in acetone, washed, and blocked with CAS-Block (Invitrogen). Staining was performed with Cy3-anti-α-smooth muscle actin (clone 1A4, Sigma) and AlexaFluor647-anti-F4/80 (clone BM8, BioLegend) diluted in CAS block, followed by washing. Sections were mounted with Abcam Fluoroshield Mounting Medium with DAPI. All images were captured with a Nikon Eclipse E800 microscope equipped with a MicroPublisher 5.0 RTV camera (QImaging) for brightfield microscopy and an EXi Blue camera (QImaging) for fluorescence microscopy using QCapture software (QImaging). Fluorescence images were merged and levelled in Adobe Photoshop.

### Leukocyte collection from tissues

Peritoneal cells were collected from body cavities by lavage. Mesenteric lymph nodes and spleens were harvested into cIMDM, homogenized through a 70 μm filter, and washed. Isolation of small intestine lamina propria cells was performed as described (*58*). In brief, 4 cm of the proximal small intestine were excised, cleaned of fecal contents, and cut into 2-3 pieces. The tissue was shaken twice in 1x HBSS (with added HEPES, 10% FBS, EDTA, and DTT) for 20 minutes to remove the epithelial layer. The tissue was then shaken at 220 rpm at 37°C for 1 hour in an Innova 4330 shaker (New Brunswick Scientific) with Collagenase IV (Sigma, C5138) in RPMI 1640 (with added 10% FBS, L-glutamine, penicillin/streptomycin, and 2-mercaptoethanol). Cells were then pelleted by centrifugation and washed.

All cells were passed through a 70 μm cell strainer before analysis. If necessary, tissues were treated with ACK lysis buffer to lyse red blood cells. Cells were counted with a hemocytometer using trypan blue.

### Flow cytometry

Cell surface staining was conducted in PBS with 0.5% BSA and 2 mM EDTA (hereafter FACS buffer) with or without 0.02% sodium azide. In brief, cells were washed in FACS buffer, blocked with α-CD16/32 (clone 2.4G2, BioXCell) for 5-10 minutes at 4°C, stained for 20 minutes at 4⁰C, and washed with FACS buffer before flow cytometry. Flow cytometry was performed on LSRFortessa X20, FACSCanto II, and LSR II instruments (all BD). Analysis was performed with FlowJo software (Treestar). Antibodies and fluorescent dyes used in this study are listed in Table S1.

Gating of cell populations was as follows (all analysis pre-gated on FSC/SSC followed by a FSC-W/FSC-A singlet gate). SILP macrophages were gated as CD45^+^F4/80^+^CD64^+^MHC-II^+^Ly6C^-^, while GMMs were gated as CD45^+^F4/80^+^CD64^+^MHC-II^lo^Ly6C^lo^. SILP eosinophils were gated as CD45^+^F4/80^+^CD64^-^CD11b^+^SSC-A^hi^, and this approach was validated in some experiments with Siglec-F staining. SILP CD3^+^ T cells were gated as CD45^+^F4/80^-^CD64^-^CD19^-^ CD3^+^. SILP CD4^+^ T cells were gated as CD19^-^F4/80^-^TCRβ^+^CD4^+^CD8^-^. LPMs were gated as CD115^+^CD11b^+^ICAM2^+^MHC-II^lo^. Peritoneal eosinophils were gated as Siglec-F^+^ICAM2^-^. Peritoneal CD4^+^ T cells were gated as ICAM2^-^TCRβ^+^CD4^+^CD8^-^. Mesenteric lymph node CD4^+^ T cells were gated as CD19^-^TCRβ^+^CD4^+^CD8^-^.

### Intracellular staining for flow cytometry

For intracellular cytokine staining of T cells, total SILP, peritoneal, and mesenteric lymph node cells were cultured at 0.5-1 million cells/well in a 96 well V-bottom non-tissue culture-treated plate for 3-4 hours at 37 °C with 8% CO2 in the presence of PMA (50 ng/ml), ionomycin (1 μM), and brefeldin A (1 μg/ml) in cIMDM. Cells were then surface stained and fixed with 4% paraformaldehyde (PFA). Cells were then washed with FACS buffer and stored overnight. To permeabilize the cells, samples were washed with 1x Perm/Wash buffer (BD, 554714) and stained for 20 minutes at 4°C, followed by washing in 1x Perm/Wash buffer and FACS buffer before flow cytometry.

For RELMα staining, the BioLegend True-Nuclear Transcription Factor Buffer set (424401) was used. After surface staining, cells were fixed with 1x Fix Concentrate buffer in Fix Diluent for 30 minutes at 4°C. Cells were then washed with FACS buffer and stored overnight. Samples were washed with 1x Perm buffer diluted in water for permeabilization. After blocking with 2% rat serum, samples were stained for 1 hour at RT with anti-RELMα antibody followed by washing in 1x Perm buffer, staining with secondary antibody for 20 min at 4°C, washing in 1x Perm buffer, and washing in FACS buffer before flow cytometry.

### Microarrays

Singlet F4/80^-^ TCRβ^+^ CD4^+^ CD8^-^ SILP cells were sorted from naïve and *H. polygyrus*-rechallenged mice using a FACSAria II (BD) into FBS, followed by lysis and RNA purification using the E.Z.N.A. MicroElute Total RNA kit (Omega Bio-Tek). RNA was submitted to the Genome Technology Access core at Washington University for cDNA synthesis (NuGen Pico SL) followed by analysis on the Affymetrix Mouse Gene 1.0 ST microarray platform. Data were analyzed with the DNASTAR ArrayStar program. All CEL files were normalized together.

Genes with an expression value of >5 (in log 2 scale) in at least one replicate were considered expressed. For generation of lists of differentially expressed genes at a ≥2-fold differential expression cutoff between groups, p-value significance of ≤0.01 by the moderated *t*-test was also required. Morpheus was used to generate heatmaps (software.broadinstitute.org/morpheus/). The Venn Diagram Plotter tool (Pacific Northwest National Laboratories, omics.pnl.gov) was used to generate Venn diagrams. Multiple differentially expressed probe sets representing a single gene were presented in heat maps, but only unique genes were counted in Venn diagrams. To assess gene set enrichment, the GSEA software from the Broad Institute was used to analyze all expressed genes for enrichment of C5 database gene sets (*59*).

### Quantification and statistical analysis

All data are from at least two independent experiments, unless otherwise indicated. Data were analyzed by unpaired two-tailed Student’s *t*-tests (Prism 7; GraphPad Software, Inc.) with p ≤ 0.05 considered significant. For relevant comparisons where no p-value is shown, the p-value was > 0.05. The NES score calculated by the GSEA software was used to account for set size effects when determining enrichment of gene sets. The GSEA-calculated FWER p value was used as a conservative measure of significance. Horizontal bars represent the mean and error bars represent the standard error of the mean (s.e.m.).

## Acknowledgements

We thank N. Zhang, B. Saunders, and G. Randolph for help with microscopy and technical advice. We thank E. Lantelme, D. Brinja, and A. Mitra for help with fluorescence activated cell-sorting. We thank J. Bando for help with flow cytometry of the gut.

## Funding

This work was supported by the National Institute of Allergy and Infectious Disease (NIAID) (AI113118 and AI132653) (B.T.E.) and a Burroughs Wellcome Fund Career Award for Medical Scientists (B.T.E.). N.N.J. was supported by grant 5T32AI007163 from the NIAID. S.C.-C.H. was supported by a Case Comprehensive Cancer Center American Cancer Society IRG Award (IRG-16-186-21) and a Cleveland Digestive Diseases Research Core Center Pilot Award (P30DK097948). M.E.C. was supported by the National Science Foundation Graduate Research Fellowship program (DGE-1745038). Research reported in this publication was supported by the Washington University Institute of Clinical and Translational Sciences grant UL1TR002345 from the National Center for Advancing Translational Sciences (NCATS) of the National Institutes of Health (NIH). The content is solely the responsibility of the authors and does not necessarily represent the official view of the NIH.

## Author contributions

N.N.J designed, performed, and analyzed experiments and wrote the manuscript. E.A.S., T.R.B., M.E.C., C.-W. L., and S.J.V.D. performed experiments. S.C.-C.H. provided reagents, protocols, and technical expertise. R.T. provided mice. T.S.S., S.J.V.D. and J.F.U. provided reagents and technical expertise. B.T.E. designed and analyzed experiments, supervised the studies, and wrote the manuscript.

## Competing interests

The authors declare no competing interests.

## Data and Materials Availability

Microarray data will be deposited in the Gene Expression Omnibus upon traditional publication.

---

**Figure S1.**
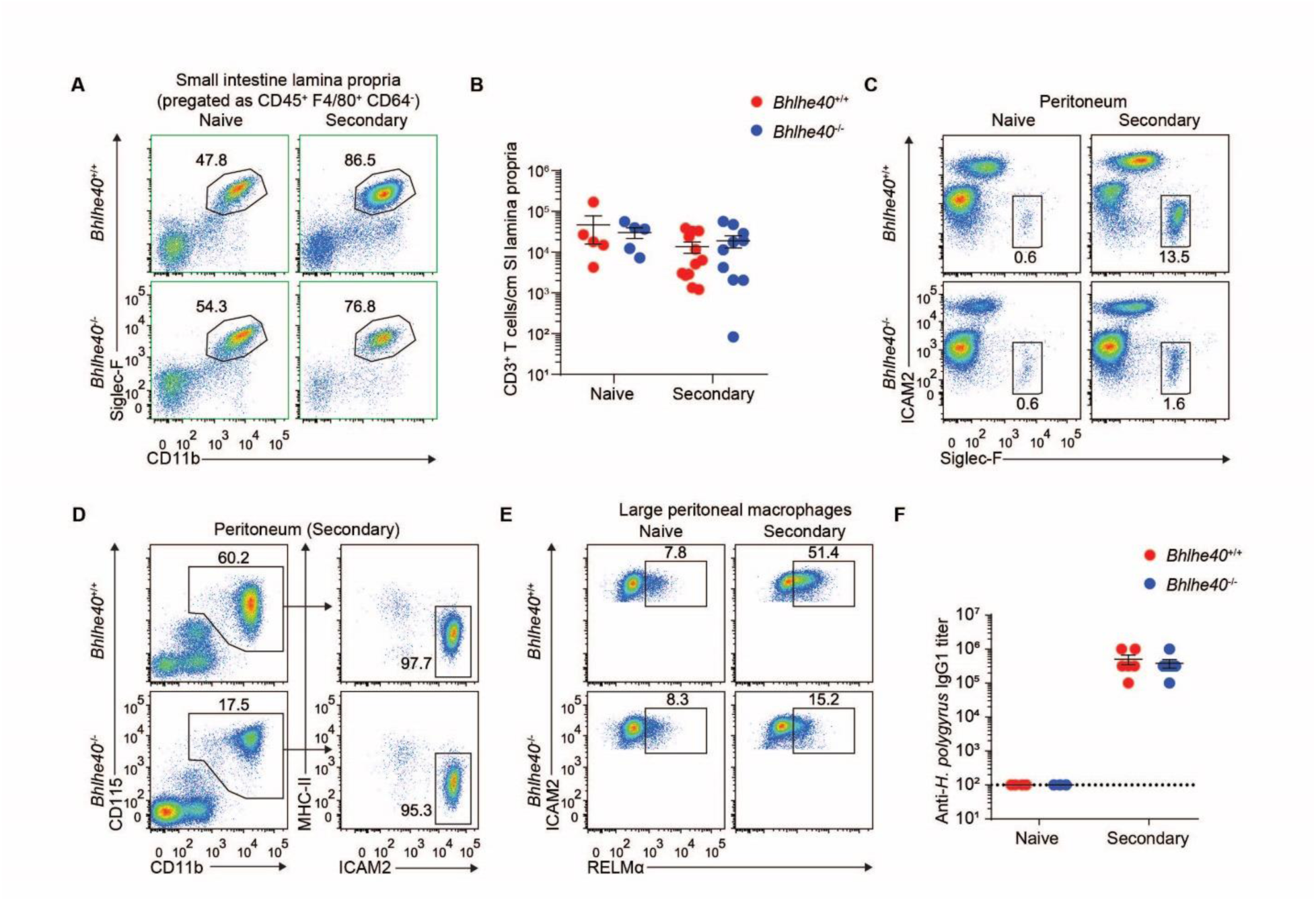
Loss of Bhlhe40 dysregulates myeloid cell responses to *H. polygyrus* rechallenge. **(A and B)** Naïve and *H. polygyrus-*rechallenged *Bhlhe40*^+/+^ and *Bhlhe40*^-/-^ mice were analyzed by flow cytometry for **(A)** SILP eosinophils and **(B)** quantitation of SILP CD3^+^ T cells. **(C-E)** Naïve and *H. polygyrus-*rechallenged *Bhlhe40*^+/+^ and *Bhlhe40*^-/-^ mice were analyzed by flow cytometry for **(C)** peritoneal eosinophils, **(D)** LPMs, and **(E)** RELMα-expressing LPMs. **(F)** Naïve and *H. polygyrus-*rechallenged *Bhlhe40*^+/+^ and *Bhlhe40*^-/-^ mice were analyzed for serum anti-*H. polygyrus* IgG1 titers. Data are representative of at least three independent experiments **(A, C-E)** or are pooled from at least 3 independent experiments **(B)** or 2 independent experiments **(F)**. Data are mean ± s.e.m. Significance calculated with an unpaired Student’s *t*-test.

**Figure S2.**
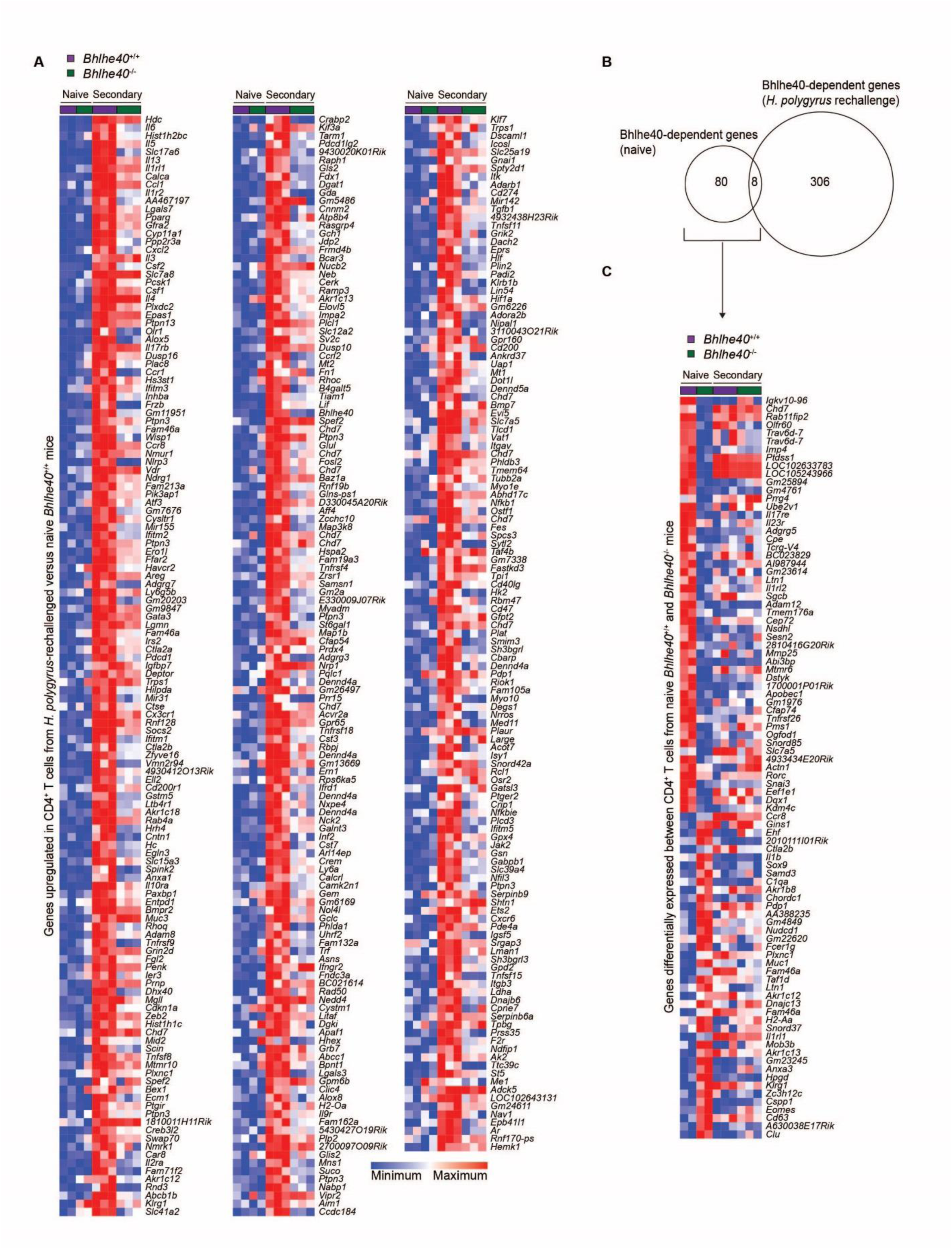
Bhlhe40 regulates distinct gene sets in CD4^+^ T cells from naïve and *H. polygyrus*-rechallenged mice. **(A)** Gene expression microarray data were analyzed for genes induced by ≥2-fold in SILP CD4^+^ T cells from *H. polygyrus*-rechallenged as compared to naïve *Bhlhe40*^+/+^ mice. **(B)** Gene expression microarray data were analyzed for shared and unique Bhlhe40-dependent genes (≥2-fold differentially expressed) in SILP CD4^+^ T cells from naïve or *H. polygyrus*-rechallenged *Bhlhe40*^+/+^ and *Bhlhe40*^-/-^ mice, depicted as a Venn diagram. **(C)** Gene expression microarray data were analyzed for genes differentially expressed by ≥2-fold in SILP CD4^+^ T cells from naïve *Bhlhe40*^+/+^ and *Bhlhe40*^-/-^ mice. Microarray data are from a single experiment.

**Figure S3.**
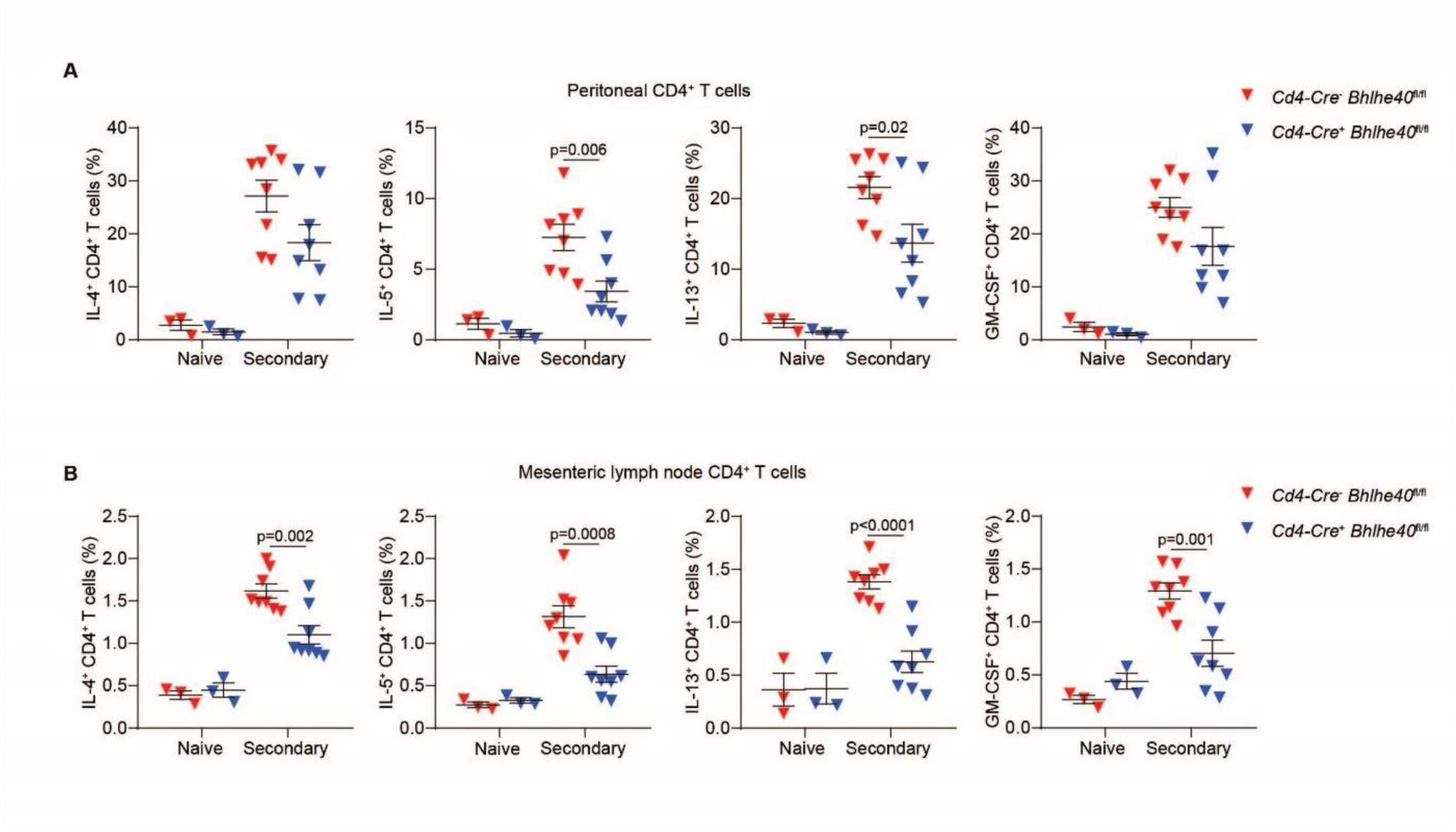
Bhlhe40 is required in T cells for normal cytokine production during *H. polygyrus* rechallenge. **(A and B)** Naïve and *H. polygyrus-*rechallenged *Cd4-Cre*^-^ *Bhlhe40*^fl/fl^ and *Cd4-Cre*^+^ *Bhlhe40*^fl/fl^ mice were analyzed by flow cytometry for quantitation of the frequency of IL-4^+^, IL-5^+^, IL-13^+^, and GM-CSF^+^ CD4^+^ T cells from the **(A)** peritoneal cavity and **(B)** mesenteric lymph nodes after *in vitro* PMA/ionomycin stimulation. Data are pooled from 2 independent experiments. Data are mean ± s.e.m. Significance calculated with an unpaired Student’s *t*-test.

**Figure S4.**
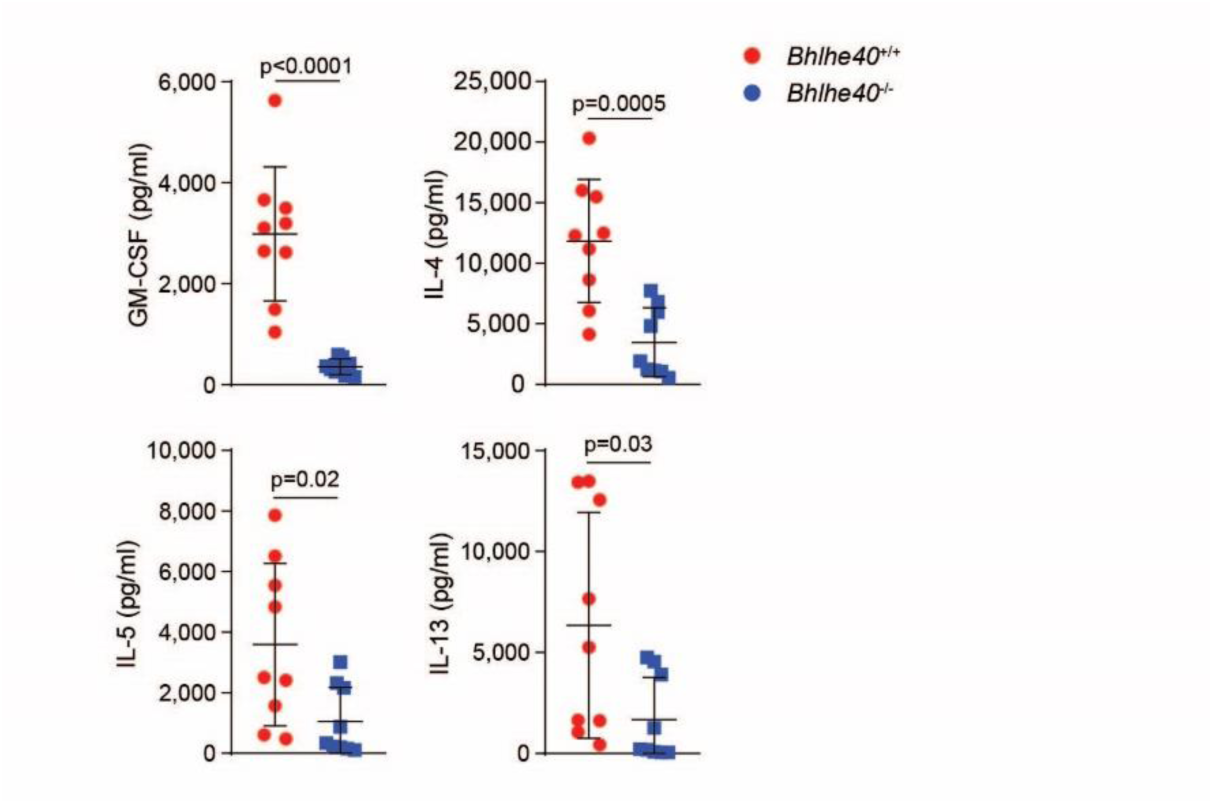
*In vitro*-polarized T_H_2 cells require Bhlhe40 for normal cytokine production. Naïve CD4^+^ T cells from *Bhlhe40*^+/+^ and *Bhlhe40*^-/-^ mice were differentiated in culture into T_H_2 cells and restimulated to assess production of GM-CSF, IL-4, IL-5, and IL-13 by ELISA. Data are pooled from 3 independent experiments. Data are mean ± s.e.m. Significance calculated with an unpaired Student’s *t*-test.

**Figure S5.**
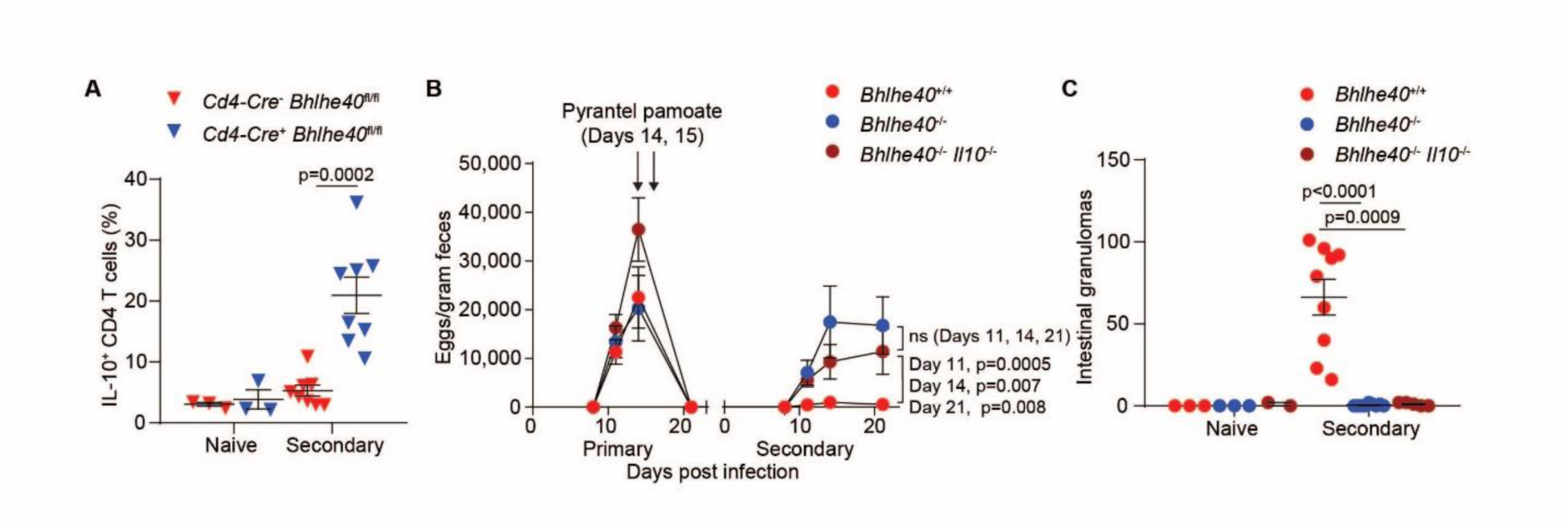
Genetic deletion of IL-10 in *Bhlhe40*^-/-^ *Il10*^-/-^ mice does not restore control of *H. polygyrus* rechallenge. **(A)** Naïve and *H. polygyrus-*rechallenged *Cd4-Cre*^-^ *Bhlhe40*^fl/fl^ and *Cd4-Cre*^+^ *Bhlhe40*^fl/fl^ mice were analyzed by flow cytometry for quantitation of the frequency of IL-10^+^ CD4^+^ T cells after *in vitro* PMA/ionomycin stimulation of SILP cells. **(B)** *H. polygyrus-*rechallenged *Bhlhe40*^+/+^, *Bhlhe40*^-/-^, and *Bhlhe40*^-/-^ *Il10*^-/-^ mice were analyzed for quantitation of *H. polygyrus* eggs/gram feces over time. **(C)** Naïve and *H. polygyrus-* rechallenged *Bhlhe40*^+/+^, *Bhlhe40*^-/-^, and *Bhlhe40*^-/-^ *Il10*^-/-^ mice were analyzed for quantitation of intestinal granulomas. Data are pooled from 2 independent experiments. Data are mean ± s.e.m. Significance calculated with an unpaired Student’s *t*-test.

**Figure S6.**
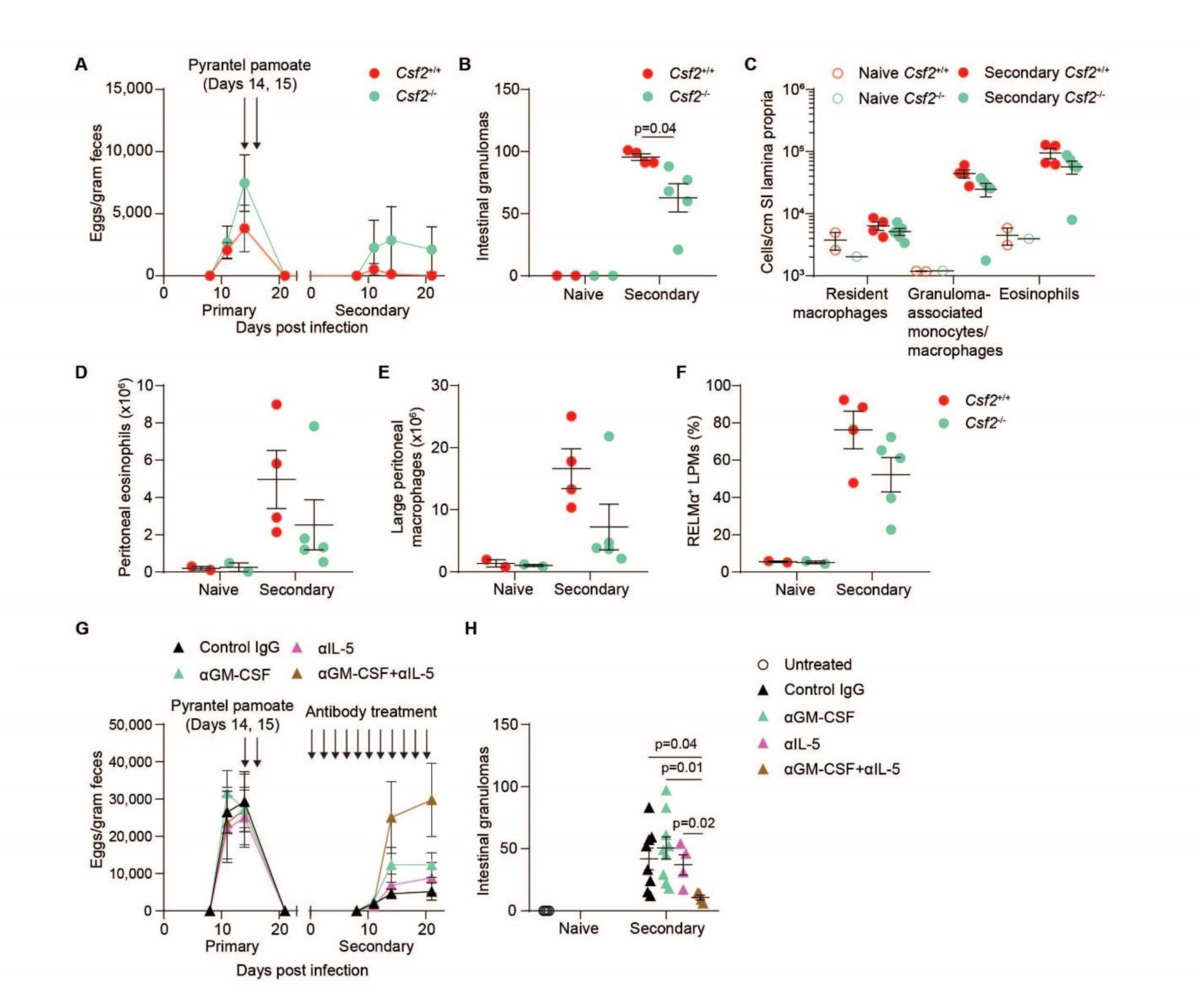
Combined deficiency in IL-5 and GM-CSF results in impaired control of *H. polygyrus* rechallenge. **(A)** *H. polygyrus-*rechallenged *Csf2*^+/+^ and *Csf2*^-/-^ mice were analyzed for quantitation of *H. polygyrus* eggs/gram feces over time. **(B)** Naïve and *H. polygyrus-* rechallenged *Csf2*^+/+^ and *Csf2*^-/-^ mice were analyzed for quantitation of intestinal granulomas. **(C)** Naïve and *H. polygyrus-*rechallenged *Csf2*^+/+^ and *Csf2*^-/-^ mice were analyzed by flow cytometry for quantitation of SILP myeloid cells. **(D-F)** Naïve and *H. polygyrus-*rechallenged *Csf2*^+/+^ and *Csf2*^-/-^ mice were analyzed by flow cytometry for quantitation of **(D)** peritoneal eosinophils, **(E)** LPMs, and **(F)** the frequency of RELMα^+^ LPMs. **(G)** Wild type mice were rechallenged with *H. polygyrus* concurrent with control IgG, αGM-CSF, αIL-5, or αGM-CSF plus αIL-5 antibody treatment and were analyzed for quantitation of *H. polygyrus* eggs/gram feces over time. **(H)** Naïve (open circles) and *H. polygyrus*-rechallenged mice (filled triangles) treated as in **(G)** were analyzed for quantitation of intestinal granulomas. Data are from single experiments. Data are mean ± s.e.m. Significance calculated with an unpaired Student’s *t*-test.

**Table S1.**
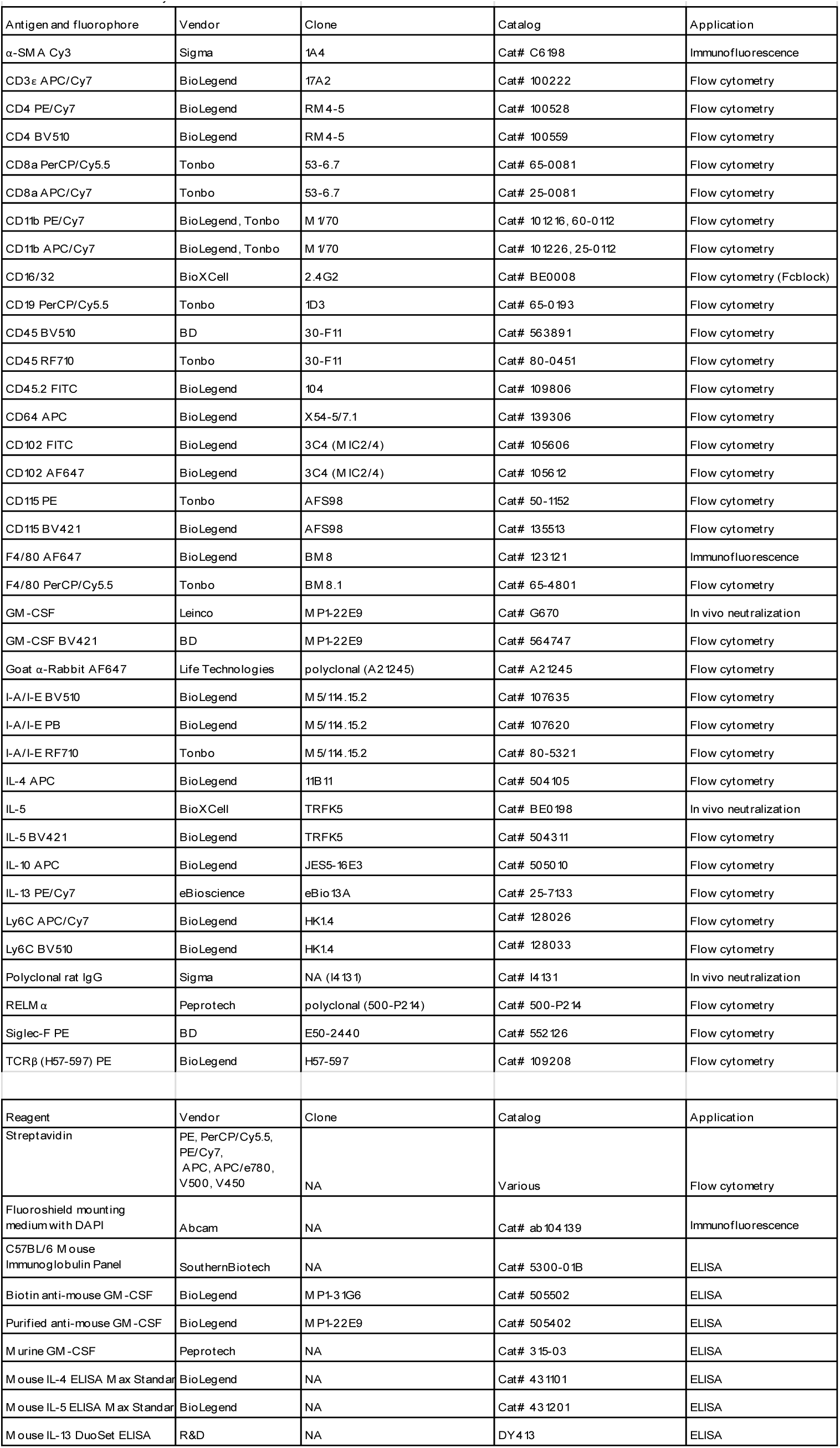
Antibodies and fluorescent dyes.

## Notes

#### Summary of Updates

This version of the manuscript has been updated to include additional data

